# Lipoprotein distribution of Apo CIII glycoforms in healthy men is linked with low TG and increased insulin sensitivity

**DOI:** 10.1101/2020.09.15.298802

**Authors:** Marina Rodríguez, Pere Rehues, Víctor Iranzo, Jorge Mora, Clara Balsells, Montse Guardiola, Josep Ribalta

## Abstract

Glycosylation of Apo CIII modulates its function in triglyceride metabolism, and some variants are associated with a protective or pro-atherogenic lipid profile. These associations have been studied in whole plasma Apo CIII proteoforms, but the proportion of Apo CIII proteoforms in individual lipoprotein fractions has been rarely evaluated. In the present study, we aim to measure the relative content of Apo CIII proteoforms in each lipoprotein fraction (VLDL, IDL, LDL and HDL) in a group of healthy subjects as a potential biomarker for triglyceride metabolism, cardiovascular risk and diabetes. Lipoprotein fractions were separated by differential ultracentrifugation of plasma samples. The relative concentrations of seven Apo CIII variants were measured by mass spectrometric immunoassay, and the complete lipoprotein profile was determined by NMR. The results showed high interindividual variability in the distribution of Apo CIII proteoforms across the study population but a uniform proportion in all lipoprotein fractions. Two Apo CIII variants, Apo CIII_0b_ and Apo CIII_1d_, were negatively correlated with plasma and VLDL triglycerides regardless of VLDL size and were associated with increased LDL size when transported in LDL particles. Apo CIII_0b_ also showed a negative correlation with lipoprotein-insulin resistance score. Therefore, Apo CIII variants can be reliably measured in lipoprotein fractions, and our results suggest that Apo CIII_0b_ and Apo CIII_1d_ have a protective role in triglyceride metabolism and insulin resistance in healthy individuals.

## INTRODUCTION

Apolipoprotein (Apo) CIII is an 8.8 kD glycoprotein that is primarily produced in the liver but also expressed in the intestine. The role of Apo CIII in triglyceride metabolism was clearly anticipated decades ago by the strong correlation observed between circulating Apo CIII and triglycerides (TG) (1). Animal models confirmed this relationship since the overexpression of the Apo CIII gene resulted in hypertriglyceridemia (2), and targeted disruption of the gene decreased TG concentrations (3). This role was recently confirmed in humans because mutations affecting the expression of the gene induced life-long reduced TG levels, which resulted in effective protection against cardiovascular disease (CVD) (4,5). Therefore, several strategies to reduce residual CVD aim to lower Apo CIII levels (6).

The proposed mechanisms by which Apo CIII contributes to dyslipidemia and cardiovascular disease are diverse and include inhibition of TG lipolysis by interacting with LPL (7,8) and stimulation of TG-rich lipoprotein secretion in liver and intestine (9). Interference with the clearance of TG-rich lipoproteins was also proposed (10), and recent data actually suggests this role as the main mechanism of action (11) because there is no *in vivo* evidence of LPL inhibition by Apo CIII under physiological conditions (12, 13). Apo CIII is also implicated in the formation of small and dense LDL particles (10), which are more atherogenic because these particles bind more efficiently to arterial proteoglycans (14). Apo CIII also enhances the binding of monocytes to the endothelium (15), and it may increase LDL susceptibility to hydrolysis by sphingomyelinases in the subendothelial space and lead to aggregation of LDL (16). In addition, new roles of Apo CIII in type 1 and type 2 diabetes were reported, including the induction of apoptosis and insulin resistance, respectively, of beta cells (17–19).

Our hypothesis is that the action of Apo CIII depends not only on the circulating concentrations of Apo CIII but also on the specific lipoprotein that carries it and the relative proportion of Apo CIII glycoforms.

The presence or absence of Apo CIII in specific lipoproteins, such as LDL and HDL, has significant clinical implications. LDL containing Apo CIII contributes to atherosclerotic CVD (20). For the atheroprotective HDL, the HDL subclass containing Apo CIII increases the risk of CVD (21). Moreover, the protective role against type 2 diabetes mellitus attributed to HDL is only observed in the fraction not containing Apo CIII (22).

Apo CIII exists in different glycoforms that originate from post-translational modifications. The most relevant forms are Apo CIII_0a_ (native form), and three variants glycosylated at Thr74: Apo CIII_0b_, with one galactose and one N-galactosamine (GalNAc); Apo CIII_1_ and Apo CIII_2_, with one and two sialic acid residues added to the Apo CIII_0b_ form, respectively (23). The influence of Apo CIII on TG metabolism varies significantly depending on the relative presence of each glycoform, with the glycoform with two molecules of sialic acid (Apo CIII_2_) having a different behavior than Apo CIII_0_ and ApoC-III_1_. This difference was observed in obese type 2 diabetic patients (23), adults with metabolic syndrome (24), in the setting of weight loss via caloric restriction (25) and in relation to LDL size (26) and triglyceride levels (27). However, all these observations were made from total plasma Apo C-III glycoforms, and not for each lipoprotein subtype.

Therefore, we hypothesized that the relative proportion of Apo CIII glycoforms could be measured in all individual lipoproteins to characterize interindividual variability and provide a useful biomarker to deepen our understanding of the role of this apolipoprotein in lipid metabolism, cardiovascular risk and diabetes.

## MATERIAL AND METHODS

### Volunteers

Twenty-four apparently healthy young men older than 18 years of age were recruited. All of them practiced physical activity and exhibited a normal lipid profile. Two of them were excluded because they were taking lipid-lowering medication.

### Biochemical analyses

Fasting venous blood samples were collected in EDTA tubes and centrifuged immediately for 15 min at 4 °C for 1500 x g to obtain the plasma. Samples were divided into aliquots and stored at −80 °C until analyses.

Standard laboratory methods were used to quantify total cholesterol, TG and HDLc. LDLc values were calculated using the Friedewald formula (28). Apolipoproteins were quantified using an immunoturbidimetric assay with specific antibodies for apolipoprotein A1 (Apo A1) (Horiba, Japan), apolipoprotein B100 (Apo B100) (Horiba, Japan) and apolipoprotein CIII (Apo CIII) (Spinreact, Spain). These analyses were adapted to the Cobas-Mira-Plus autoanalyzer (Roche Diagnosis, Spain).

### Lipoprotein separation

Lipoprotein fractions were separated via sequential ultracentrifugation after progressively increasing the solvent density (29) using the ultracentrifuge model Optima XPN-100 (Beckman Coulter, California, USA) and a Kontron 45.6 fixed-angle rotor at 37000 rpm and 4 °C for different time intervals: 16 h for chylomicrons (QM) and VLDL, 20 h for IDL, 20 h for LDL and 40 h for HDL. To separate these fractions, density solutions of 1.006, 1.019, 1.063 and 1.239 g/ml were used, respectively.

### Apo CIII proteoforms using mass spectrometric immunoassay (MSIA)

Analysis of Apo CIII was performed using immunoprecipitation of Apo CIII complexes coupled to a mass spectrometric analysis of the intact Apo CIII protein, as previously described (30). Briefly, affinity columns were prepared by immobilizing the biotinylated antibody (2.5 μg of anti-Apo CIII) with streptavidin magnetic beads (Dynabeads, Life Technologies, Grand Island, NY). Following sample preparation, a total of 120 μl of plasma sample or lipoprotein subfractions diluted 120-fold in PBS, 0.1 % Tween were incubated for 45 min with magnetic beads coated with a streptavidin-biotin anti-Apo CIII antibody to capture Apo CIII proteoforms from each analytical sample. After rinsing the nonspecifically bound proteins, captured apolipoproteins were eluted directly onto a 96-well formatted MALDI target using a Sinapinic acid matrix. Linear mass spectra were acquired from each sample spot using an UltrafleXtrem III MALDI-TOF/TOF instrument (Bruker, Germany) operating in positive ion mode. An average of 3000 laser shot mass spectra were saved for each sample spot. Mass spectra were internally calibrated using protein calibration standard-I and further processed using Flex Analysis 3.0 software (Bruker Daltonics). All peaks representing Apo CIII and their proteoforms were integrated. To assess the consistency of the ionization efficiency and reproducibility between and within runs, a quality control plasma sample was run in triplicate with each analysis.

For the sialylated proteoforms of Apo CIII, seven main Apo CIII proteoforms were detected (Suppl Figure 1): Apo CIII_0a_ is the native Apo CIII without glycosylation; C-III_0b_ is glycosylated with one galactose (Gal) and one N-acetylgalactosamine (GalNAc); C-III_0f_ is a more glycosylated proteoform with two molecules of galactose, two of N-acetylgalactosamine and three molecules of fucose((Gal)2(GalNAc)2(Fuc)3); C-III_1_ has the same glycosylation as C-III_0b_ and one molecule of sialic acid ((Gal)1(GalNAc)1(NeuAc)1); C-III_1d_ is C-III_1_ but without one alanine; C-III_2_ is the same as C-III_1_ but with two molecules of sialic acid ((Gal)1(GalNAc)1(NeuAc)2); and C-III_2d_ is C-III_2_ but without one alanine.

### Liposcale® Test: Nuclear magnetic resonance (NMR) lipoprotein profile

Total plasma lipids and distributions of lipoprotein subclasses were analyzed via NMR spectroscopy by Biosfer Teslab (Reus, Spain). The Liposcale® test is a novel advanced lipoprotein test based on 2D diffusion-ordered ^1^H-NMR spectroscopy and uses diffusion coefficients (31). This technique provides the particle size and number of the main types of lipoprotein (VLDL, LDL, and HDL), dividing each one in large, medium and small sizes according to increases in molecular weight. ^1^H-NMR approaches are based on regression methods, which also allow for the determination of the cholesterol and triglyceride concentrations of lipoproteins (32). The ^1^H-NMR was carried out on EDTA plasma stored and thawed just prior to the analysis.

### LP-IR score

The Lipoprotein Insulin Resistance (LP-IR) score was calculated as described by Shalaurova et al. (33). Briefly, six parameters of lipoproteins (VLDL size, large VLDL particles, LDL size, small LDL particles, HDL size and large HDL particles) were divided into several categories, and a value was given to each category. The six values were added to obtain a score between 0 and 100 to indicate insulin resistance.

### Statistical analyses

The normal distribution of variables was assessed using the Shapiro-Wilk test. Partial correlations between Apo CIII proteoforms and lipid and lipoprotein profile were adjusted for age, BMI, and total Apo CIII or TG levels.

Statistical significance was set at *p* < 0.05, and all statistical analyses were evaluated using SPSS Statistical Software version 23 (SPSS Inc., Chicago, USA).

## RESULTS

### Description of the study population

We studied 22 young healthy males. Anthropometric and conventional lipid profile data showed that all parameters were within the normality range (Table 1). The detailed lipid profiles determined by the Liposcale® test using nuclear magnetic resonance were also carried out. Data are shown in Suppl Table 1.

**Table 1.** Characteristics of the study population.

|  |  |
| --- | --- |
| **Age, years** | **25.5 ± 5.1** |
| **Body Mass Index, kg/m<sup>2</sup>** | **23.3 ± 2.4** |
| **Cholesterol, mg/dl** | **170.5 ± 22.2** |
| **LDLc, mg/dl** | **92.1 ± 23.2** |
| **HDLc, mg/dl** | **58.5 (52.8 – 70.8)** |
| **Triglycerides, mg/dl** | **79.6 ± 19.2** |
| **Apo A1, mg/dl** | **142.0 ± 17.8** |
| **Apo B100, mg/dl** | **79.0 ± 17.2** |
| **Apo CIII, mg/dl** | **7.6 ± 2.8** |

Values of variables normally distributed are represented as the means ± SD, and variables not normally distributed as medians (IQR).

### Description of proteoforms and distribution

We measured the seven main Apo CIII proteoforms in all lipoproteins in 22 healthy men, except for the Apo CIII_0f_ proteoform, which was not detected in some lipoprotein fractions of 4 individuals because it is a minority proteoform. Figure 1 shows the mean values of the average relative distribution of each proteoform in plasma, VLDL, IDL, LDL and HDL. When all subjects were studied together, the relative distribution of all seven proteoforms was constant in plasma and the four isolated lipoproteins, with Apo CIII_1_ as the most abundant form, followed by Apo CIII_2_ and Apo CIII_0b_.

**Figure 1.**
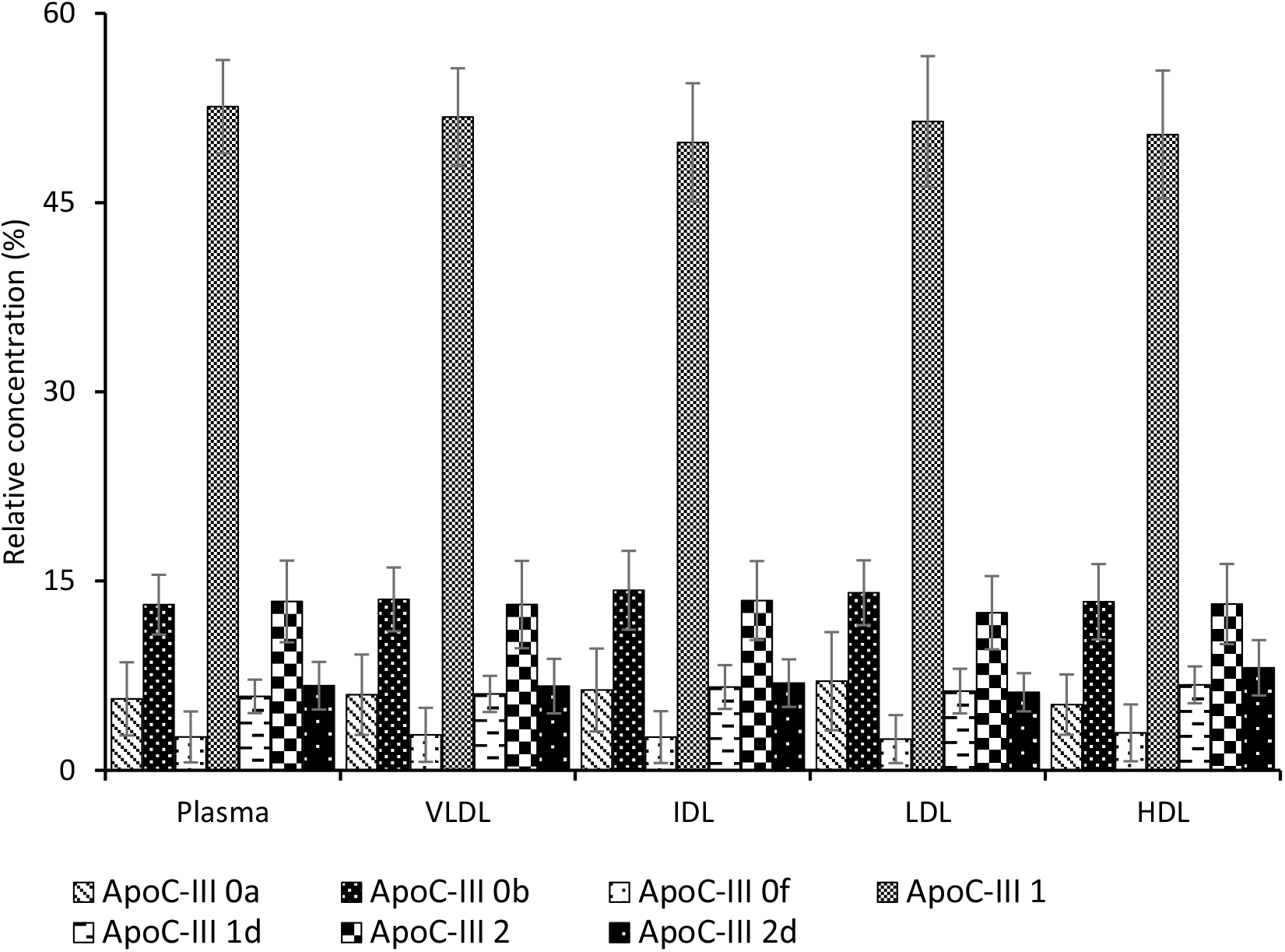
Distribution of all different proteoforms of Apo CIII in plasma and lipoproteins in healthy men. Means of the relative concentration of Apo CIII proteoforms in each lipoprotein fraction and SD are shown. The relative concentration of each proteoform is constant in all lipoprotein fractions and plasma. Apo CIII_1_ is the most abundant proteoform, followed by Apo CIII_0b_ and Apo CIII_2_; then Apo CIII_0a_, Apo CIII_1d_ and Apo CIII_2d_; and finally Apo CIII_0f_, the least abundant of all proteoforms analyzed.

Suppl Figure 2 shows the relative distribution in plasma and lipoproteins for each individual proteoform in each of the 22 study participants. There was high interindividual variability in the relative distribution of each Apo CIII proteoform, but, for each subject, the proportion of each proteoform was maintained in all lipoproteins. Hierarchical clustering revealed the existence of 4 subgroups in the study population according to the distribution of Apo CIII glycoforms (Figure 2). Subgroup 1 was composed of subjects with a combination of high C-III_2_ and low C-III_0_ glycoforms. In contrast, subgroup 2 was composed of subjects with low C-III_2_ and high C-III_0_ glycoforms. In subgroups 1 and 2, half of the subjects had high C-III_1_ (1a and 2a) and half had low C-III_1_ (1b and 2b).

**Figure 2.**
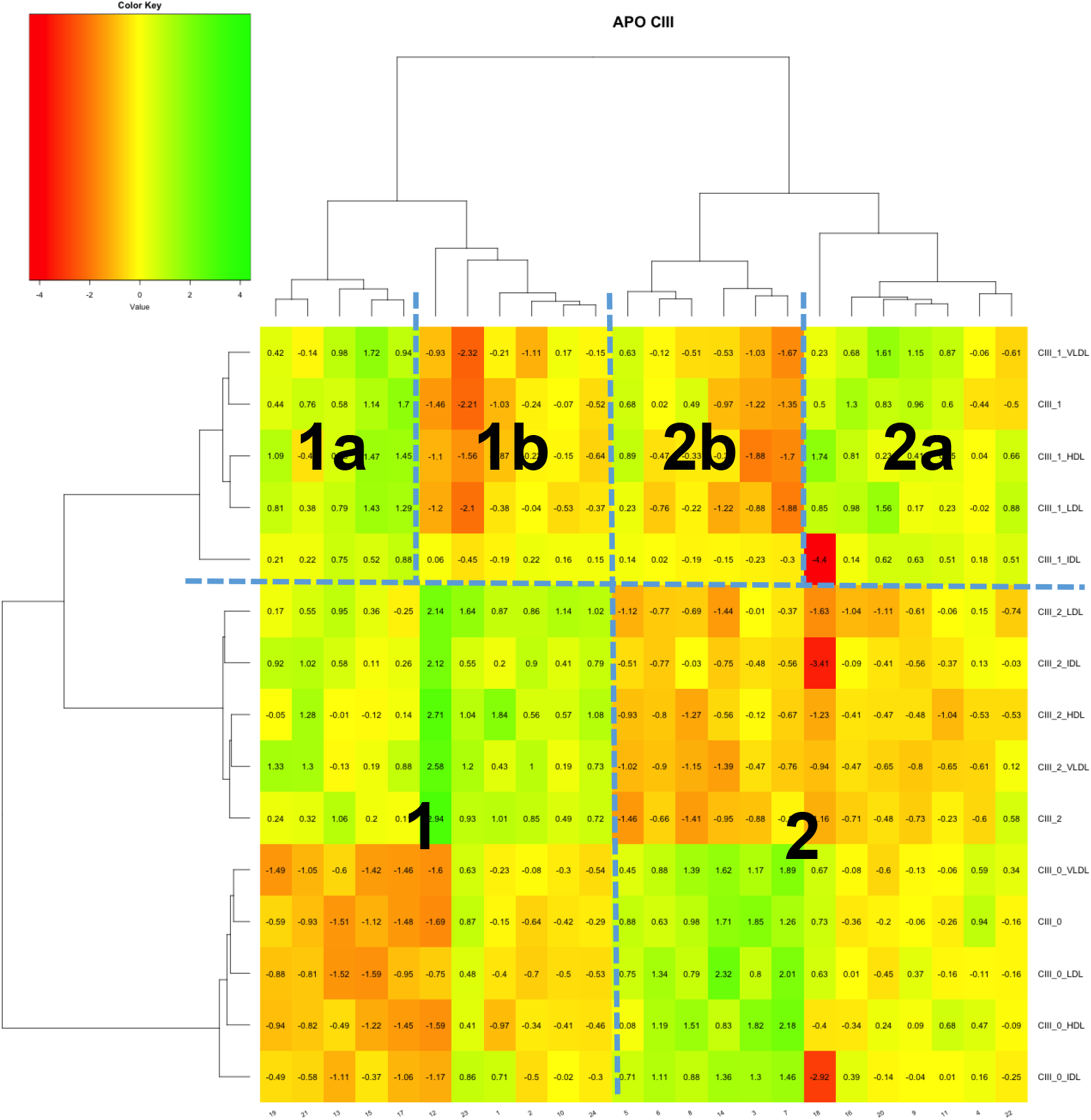
Hierarchical clustering of Apo CIII glycoforms. Study subjects can be divided in two subgroups according to their distribution of Apo CIII proteoforms. Subgroup 1 consists of subjects with high Apo CIII_2_ and low Apo CIII_0_ while subgroup 2 consists of subjects with low Apo CIII_2_ and high Apo CIII_0_. Each subgroup is further divided according to the relative concentration of Apo CIII_1_ in *a* (high) and *b* (low).

### Relationship of Apo CIII proteoforms, TG and triglyceride-rich lipoproteins (TRL)

Because Apo CIII has an obvious role in TG metabolism we focused our initial analyses on plasma TG and VLDL and explored the association between Apo CIII proteoforms, TG and TG-rich lipoproteins.

Apo C-III_0b_ and Apo C-III_1d_ proteoforms were the only forms that correlated significantly with triglycerides. The relative concentration of plasma ApoC-III_0b_ showed a strong and negative correlation with circulating plasma triglycerides (r=−0.855, p<0.001) and VLDL triglycerides (r=−0.899, p<0.001), independent of total Apo CIII (Table 2).

**Table 2.**
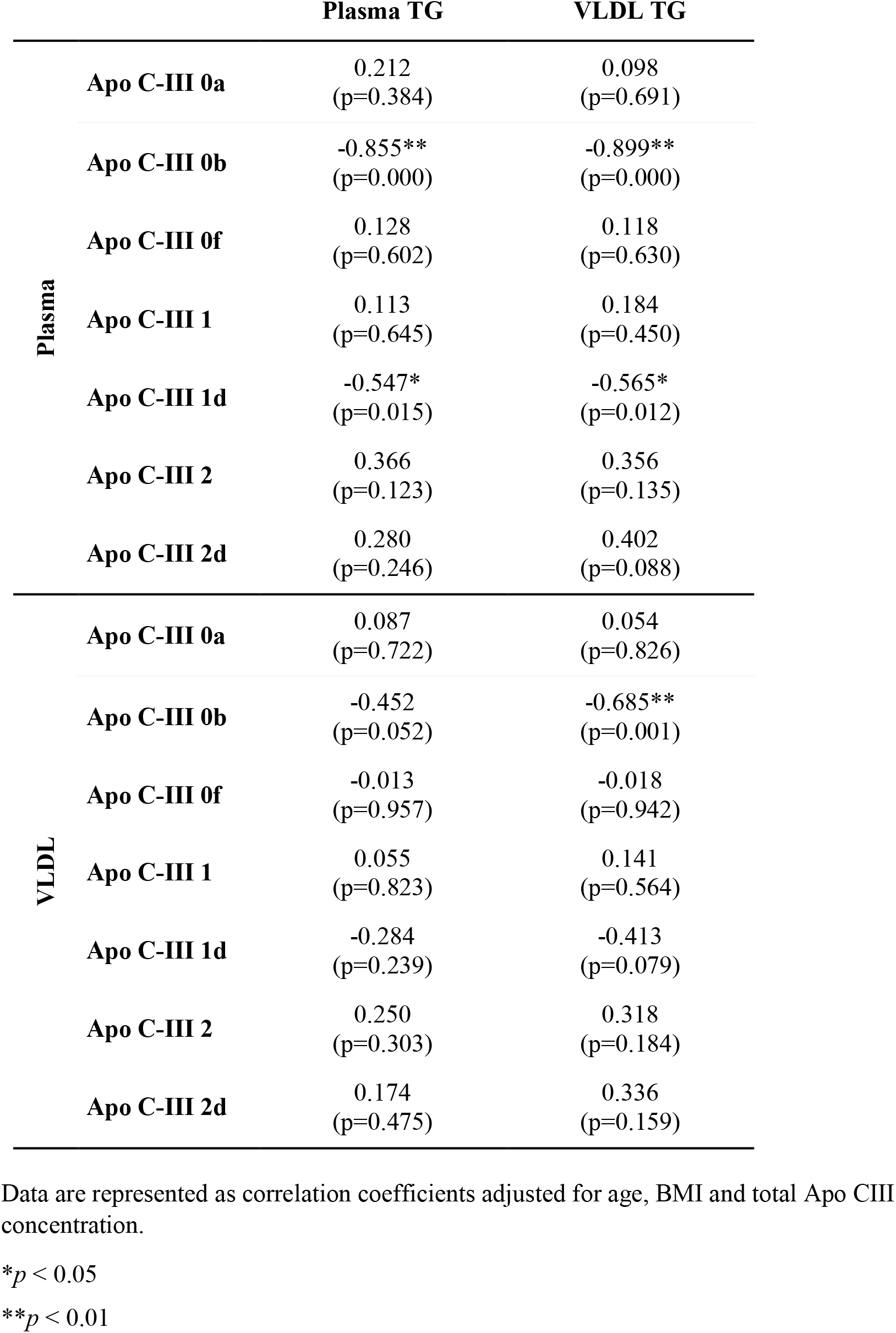
Correlation between all Apo CIII proteoforms and TG in plasma and VLDL.

The ApoC-III_0b_ proteoform carried in VLDL particles showed similar correlations with plasma TG and VLDL-TG levels (r=−0.452, p=0.052 and r=−0.685, p=0.001, respectively) controlling for total ApoC-III concentration (Table 2). This correlation was maintained in all VLDL lipoprotein sizes (large, medium and small; Table 3).

**Table 3.** Correlation between all Apo CIII proteoforms measured in VLDL and VLDL characteristics.

|  | VLDL particles |  |  |  | Size |
| --- | --- | --- | --- | --- | --- |
|  | Total | Large | Medium | Small |  |
| **VLDL** |  |  |  |  |  |
| **CIII-0a** | 0.077<br>(p=0.755) | 0.099<br>(p=0.687) | 0.077<br>(p=0.754) | 0.074<br>(p=0.764) | 0.101<br>(p=0.680) |
| **CIII-0b** | -0.655**<br>(p=0.002) | -0.582**<br>(p=0.009) | -0.646**<br>(p=0.003) | -0.648**<br>(p=0.003) | -0.282<br>(p=0.242) |
| **CIII-0f** | -0.022<br>(p=0.929) | 0.007<br>(p=0.979) | 0.029<br>(p=0.908) | -0.034<br>(p=0.889) | 0.210<br>(p=0.388) |
| **CIII-1** | 0.149<br>(p=0.544) | -0.026<br>(p=0.915) | 0.037<br>(p=0.881) | 0.181<br>(p=0.458) | -0.315<br>(p=0.189) |
| **CIII-1d** | -0.401<br>(p=0.089) | -0.360<br>(p=0.130) | -0.381<br>(p=0.108) | -0.400<br>(p=0.089) | -0.148<br>(p=0.545) |
| **CIII-2** | 0.266<br>(p=0.271) | 0.391<br>(p=0.098) | 0.370<br>(p=0.119) | 0.230<br>(p=0.343) | 0.426<br>(p=0.069) |
| **CIII-2d** | 0.330<br>(p=0.168) | 0.294<br>(p=0.221) | 0.306<br>(p=0.203) | 0.331<br>(p=0.167) | 0.049<br>(p=0.841) |

Data are represented as correlation coefficients adjusted for age, BMI and Apo CIII.
\*\* $p < 0.01$

Similar results were observed for relative concentrations of plasma Apo CIII_1d_ with plasma TG (r=−0.547, p=0.015) and VLDL-TG (r=−0.565, p=0.012), adjusting for total Apo CIII concentration. The correlations observed for Apo CIII_1d_ carried in VLDL particles followed similar tendencies but with lower coefficients and no significance (Table 2). Therefore, higher relative contents of ApoC-III_0b_ and ApoC-III_1d_ were associated with lower TG levels.

Although the other Apo CIII proteoforms did not correlate significantly with TG, Apo CIII_2_, Apo CIII_2d_ and plasma Apo CIII_0a_ tended to have a positive correlation with TG (Table 2).

### Relationship of ApoC-III proteoforms with the complete lipoprotein profile

We also explored other potential associations between all seven Apo CIII proteoforms and detailed lipid parameters, such as LDL and HDL lipoproteins or the main apolipoproteins.

The abovementioned Apo CIII_0b_ proteoform also correlated positively with HDLc (r=0.571, p=0.008), ApoA1 (r=0.495, p=0.026), and the amount of total, medium and small HDL particles (r=0.543, p=0.013; r=0.592, p=0.006; r=0.469, p=0.037; respectively), while a negative correlation with VLDL cholesterol was observed (r=−0.733, p<0.001). The ApoC-III_0b_ transported in LDL particles also correlated positively with LDL particle size (r=0.648, p=0.003; independent of total Apo CIII levels) (Table 4).

**Table 4.** Correlation between all Apo CIII proteoforms in LDL and LDL size.

|  |  | LDL size |
| --- | --- | --- |
| LDL | Apo C-III 0a | -0.413<br>(p=0.078) |
|  | Apo C-III 0b | 0.648**<br>(p=0.003) |
|  | Apo C-III 0f | 0.048<br>(p=0.847) |
|  | Apo C-III 1 | -0.449<br>(p=0.054) |
|  | Apo C-III 1d | 0.529*<br>(p=0.020) |
|  | Apo C-III 2 | 0.289<br>(p=0.230) |
|  | Apo C-III 2d | 0.339<br>(p=0.155) |

Data are represented as correlation coefficients adjusted for age, BMI and total Apo CIII.
\* $p < 0.05$
\*\* $p < 0.01$

However, some of these correlations (ApoA1, small and large HDL particle numbers) lost statistical significance after adjustment for ApoC-III levels, and all of the correlations except HDL cholesterol and LDL size were lost after adjusting for TG in partial correlations, which highlights the significant role of TG in the observed associations.

ApoC-III_0a_ correlated positively with total cholesterol (r=0.480, p=0.032) and IDL cholesterol (r=0.481, p=0.032), but only IDL cholesterol was significant after adjusting for total Apo CIII levels. None of the factors were significant after adjusting for TG.

ApoC-III_1_ correlated positively with the amount of large HDL particles (r=0.464, p=0.039).

ApoC-III_1d_ correlated negatively with VLDL cholesterol (r=0.532, p=0.016) and, when transported in LDL particles, it also correlated with LDL particle size (r=0.529, p=0.020; Table 4). These correlations remained significant after adjusting for total Apo CIII, but significance was lost when adjusting for TG;only LDL size remained significant.

Finally, a negative correlation was observed between Apo CIII_2_ and small HDL particles (r=−0.446, p=0.049), which was no longer significant after adjusting for total Apo CIII or TG. A positive correlation between Apo CIII_2d_ and VLDL large particle number (r=0.451, p=0.046) was observed, which was maintained after controlling for TG or total Apo CIII levels.

### LP-IR score

An insulin resistance score based on alterations of lipoprotein characteristics was recently proposed as a novel, BMI-independent test for insulin resistance (33). We calculated the lipoprotein-insulin resistance (LP-IR) score for all of the individuals in the study and found a negative association with the relative content of Apo CIII_0b_ in plasma (r=−0.617, p= 0.004; Figure 3) and in the IDL (r=−0.577, p= 0.010), LDL (r=0.454, p=0.045) and HDL (r=-0.541, p=0.014) fractions. These correlations were independent of Apo CIII total concentration but lost significance after adjusting for TG.

**Figure 3.**
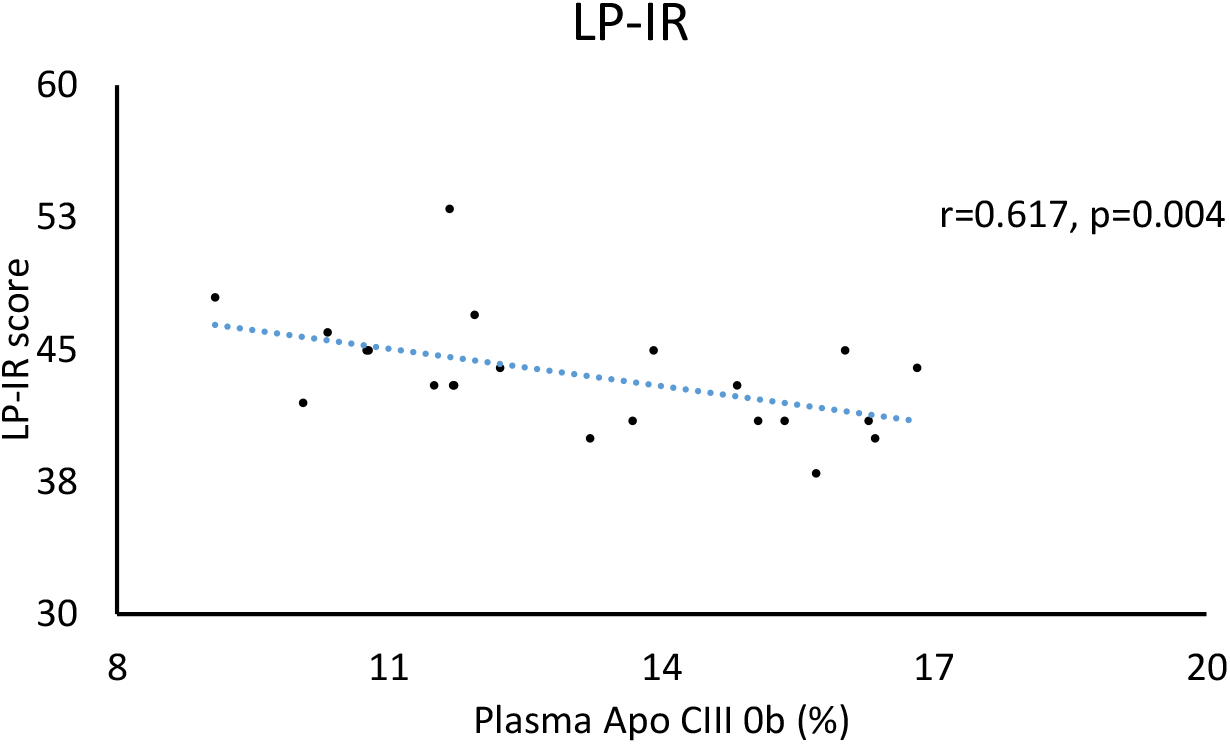
Correlation between Apo CIII_0b_ proteoform in plasma and LP-IR score. The relative concentration of plasma Apo CIII_0b_ correlated negatively with the LP-IR score, a scale between 0 and 100 that indicates insulin resistance, showing a protective role of Apo CIII_0b_ proteoform in insulin resistance.

## DISCUSSION

There is convincing evidence that life-long reduced TG levels due to APOC3 mutations result in effective protection against cardiovascular disease (4,5), and Apo CIII is also linked to the onset of type 1 and 2 diabetes mellitus. Therefore, Apo CIII is a very promising biomarker to study the cardiovascular risk associated with diabetes. Understanding the mechanism of action of Apo CIII is complex because scientific evidence indicates that the effects of Apo CIII depend on the lipoprotein to which it is bound (20,21) and the relative abundance of the Apo CIII glycoforms (23–26). Only the non-glycated, mono-sialo and di-sialo forms were measured in VLDL, LDL and HDL (24); and most of the published studies only considered total Apo CIII glycoforms and not its distribution in individual lipoproteins.

Because of its potential as a biomarker, we measured seven Apo CIII glycoforms in plasma and each of the individual lipoproteins (VLDL, IDL, LDL and HDL) and characterized their interindividual variability.

We demonstrated that all seven Apo CIII glycoforms can be reliably measured in plasma and each separate lipoprotein (VLDL, IDL, LDL and HDL) in young healthy subjects. Based on the obtained measurements, we made two main observations. There was significant interindividual variability in the relative distribution of the seven glycoforms but, in each given subject, the distribution was maintained throughout the different lipoproteins. Consistent with several previous observations, the correlations with TG and lipoproteins differ depending on the Apo CIII glycoform.

The fact that Apo CIII glycoform distribution was maintained across the different lipoproteins naturally raises the question of the need to measure Apo CIII glycoform distribution in each lipoprotein when plasma may be an equally valid indicator. Although this is one possible scenario, another possibility is that this distribution is a characteristic of normal metabolism, which may be disrupted in the presence of metabolic alterations, being such disruption itself a useful biomarker. Obviously, this hypothesis deserves further studies in diabetic, obese or metabolic syndrome patients. To set up the measurement of Apo CIII in individual lipoproteins, we chose a population of healthy young males because we assumed that the ability to detect the seven glycoforms in low-concentration lipoproteins and low Apo CIII concentrations would be an indicator of the robustness of the method.

Our results show that Apo CIII_0b_ and Apo CIII_1d_ proteoforms are negatively associated with TG and TRL levels in healthy young men and remained associated after adjusting for total Apo C-III concentrations. However, no other significant correlations were found among the analyzed proteoforms. Our results are consistent with those reported by Olivieri et al. in a cohort of CAD patients where a negative tendency was reported between Apo CIII_0_ proteoforms and TG (34). However, there is controversy because in previous studies with prediabetic and diabetic patients, baseline levels of Apo CIII_0b_ correlated positively with plasma TG, and the ratio Apo CIII_2_/Apo CIII_1_ showed an inverse correlation with plasma TG (27). The same results were reported in overweight and obese patients (23, 26), in whom all proteoforms except Apo CIII_2_ positively correlated with TG levels.

The atheroprotective association of 0b and 1d in our study was also reinforced by the correlations with large LDL, which is certainly a direct consequence of the effect on TG levels, and suggests a broad effect of Apo CIII proteoforms in lipid metabolism. The results in the literature about Apo CIII proteoforms and LDL size are more scarce and less clear. The proportion of Apo CIII_0b_ was associated with the LDL4 subtype (small) in prediabetic and diabetic patients (27), and the absolute concentration of Apo CIII_0b_ inversely correlated with LDL size in the study by Mendoza et al. (26), although when normalized to the Apo CIII_0a_ proteoform the correlation became positive.

The differences observed between our results and previous reports may be due to the characteristics of our group of study, which was very different from previous studies because they exhibited no cardiovascular-related issues and were not receiving any lipid-lowering therapy. In addition, our analyses were made on the relative content of Apo CIII glycoforms (percentage of one glycoform over the seven measured glycoforms), and most of the previous results used absolute concentrations or ratios between two glycoforms.

Because many functions of Apo CIII in lipid metabolism were described, one possible explanation for the observed correlations is that an increased proportion of the 0b glycoform would affect the inhibitory role of Apo CIII on the uptake of TRLs. TRL receptors have different affinities for lipoproteins depending on which Apo CIII glycoforms are associated with the particle (35). Apo CIII may interfere with the Apo E-mediated hepatic uptake of TRL (11, 36), and the 0b glycoform, with a smaller glycan moiety, may exert a decreased interference on Apo E binding to the LDLR or LRP than the other glycoforms. Therefore, a higher relative amount of Apo CIII_0b_ would allow for a more efficient uptake and lower plasma concentrations of TG and VLDL particles. Another possible mechanism by which Apo CIII_0b_ may be associated with lower TG levels is that Apo CIII_0b_-containing TRL may be cleared by receptors other than LDLR or LPR, such as HSPG or ASGPr (the latter recognizes exposed Gal and GalNac residues). However, none of these associations have been described, and further research must be done to explore these hypotheses.

In type 1 and type 2 diabetes, increased levels of Apo CIII promote Ca^2+^ channel activity and the apoptosis of beta cells or impairment of beta cell functions (18). This role of Apo CIII may also depend on its glycosylation state because increased levels of the sialylated forms and decreased levels of the nonsialylated forms were observed in type 1 diabetes patients compared to healthy controls (37). We explored the associations of glycoforms with the insulin resistance marker LP-IR score and found a negative correlation with Apo CIII_0b_ proteoform. This result is consistent with the results observed in TRL metabolism, which suggest that some Apo CIII functions are glycoform-dependent, and supports the protective role of Apo CIII_0b_ also in insulin resistance. However, further investigation in this field is needed to disclose the role of glycosylation on the process of insulin resistance.

In conclusion, the present study shows that Apo CIII proteoforms can be measured in all lipoprotein fractions of healthy subjects and present different interindividual distributions that may be evaluated as a potential biomarker. Among the seven proteoforms analyzed, an atheroprotective role of Apo CIII_0b_ and Apo CIII_1d_ on TG metabolism is proposed. Consistently, the association of Apo CIII_0b_ with the LP-IR score suggests that the protective role of this proteoform also affects insulin resistance. Therefore, a higher relative content of these proteoforms may attenuate the detrimental effects of Apo CIII, but further research is necessary to clarify the role of the different Apo CIII proteoforms on TG metabolism and insulin resistance.

## DATA AVAILABILITY

The data analyzed in this study are available from the corresponding author J. R. upon request.

## ACKNOWLEDGEMENTS

This work was supported by the Spanish Ministerio de Economía y Competitividad (PI20/00514), Fondo Europeo de Desarrollo Regional (FEDER), and CIBERDEM (CIBER de Diabetes y Enfermedades Metabólicas Asociadas), which are initiatives of ISCIII (Instituto de Salud Carlos III). P. R. is a recipient of a predoctoral fellowship from the Ministerio de Universidades (grant number  FPU19/04610).

